# Epigenetic conflict on a degenerating Y chromosome increases mutational burden in Drosophila males

**DOI:** 10.1101/2020.07.19.210948

**Authors:** Kevin H.-C. Wei, Lauren Gibilisco, Doris Bachtrog

## Abstract

Large portions of eukaryotic genomes consist of transposable elements (TEs), and the establishment of transcription-repressing heterochromatin during early development safeguards genome integrity in Drosophila. Repeat-rich Y chromosomes can act as reservoirs for TEs (‘toxic’ Y effect), and incomplete epigenomic defenses during early development can lead to deleterious TE mobilization. Here, we contrast the dynamics of early TE activation in two Drosophila species with vastly different Y chromosomes of different age. Zygotic TE expression is elevated in male embryos relative to females in both species, mostly due to expression of Y-linked TEs. Interestingly, male-biased TE misexpression diminishes across development in *D. pseudoobscura*, but remains elevated in *D. miranda*, the species with the younger and larger Y chromosome. The repeat-rich Y of *D. miranda* still contains many actively transcribed genes, which compromise the formation of silencing heterochromatin. Elevated TE expression results in more *de novo* insertions of repeats in males compared to females. This lends support to the idea that the ‘toxic’ Y chromosome can create a mutational burden in males when genome-wide defense mechanisms are compromised, and suggests a previously unappreciated epigenetic conflict on evolving Y chromosomes between transcription of essential genes and silencing of selfish DNA.

## Introduction

In most animals, the zygotic genome is initially inactive and epigenetically ‘naïve’ (i.e. it contains few chromatin features) (Li *et al*. 2014), and the first stages of embryonic development are solely controlled by maternal proteins and transcripts (Newport and Kirschner 1982a; b). Concordant with genome-wide activation of zygotic expression, the embryo also initiates silencing of genomic regions whose transcription would be harmful (Haig 2016). In particular, large fractions of eukaryotic genomes consist of repetitive DNA including transposable elements (TEs) (Padeken *et al*. 2015), and transcriptional activation of TEs could result in their mobilization, causing insertional mutations and genomic instability (Hedges and Deininger 2007). Silencing of repeats is achieved in part through establishment of constitutive heterochromatin in all cells during early development at repetitive DNA at centromeres, telomeres, and along the repeat-rich Y chromosome (Girton and Johansen 2008; Elgin and Reuter 2013). TEs are in a constant evolutionary battle with their host genome to avoid silencing, and may have evolved mechanisms to mobilize early in embryogenesis before widespread heterochromatin is in place (Bourque *et al*. 2018). In many species, including Drosophila, the Y chromosome consists almost entirely of repetitive DNA, and male embryos may thus be especially challenged to silence their TEs (Brown *et al*. 2020a).

Here, we contrast TE activation and heterochromatin formation during early embryogenesis in two closely related Drosophila species (**Fig. 1**). *Drosophila pseudoobscura* and *D. miranda* diverged only 3Ma (million years ago) and show a similar repeat complement, with most TEs being shared between species (Hill and Betancourt 2018; Bracewell *et al*. 2019). Yet, the size of their Y chromosome differs dramatically: A fusion between an autosome and the ancestral Y created a neo-Y in *D. miranda* about 1.5Ma (Bachtrog and Charlesworth 2002) (**Fig. 1A**), which has dramatically expanded in size (from ∼25Mb to almost 100Mb; (Mahajan *et al*. 2018)). This drastic size expansion is driven almost entirely by the accumulation of TEs on the neo-Y (Mahajan *et al*. 2018); for example, the most common TE on the neo-Y (the ISY element) is inserted roughly 22,000 times on the neo-Y (and occupies over 16Mb of sequence), while it only shows about 1,500 copies on the neo-X (<1Mb of sequence)(Mahajan *et al*. 2018). High-quality genome assemblies that contain large amounts of highly repetitive DNA, including pericentromeric regions and Y-linked sequences, exist for both species (Mahajan *et al*. 2018; Bracewell *et al*. 2019), which allow us to study TE expression and heterochromatin formation at the molecular level, and the role of the Y chromosome.

**Figure 1.**
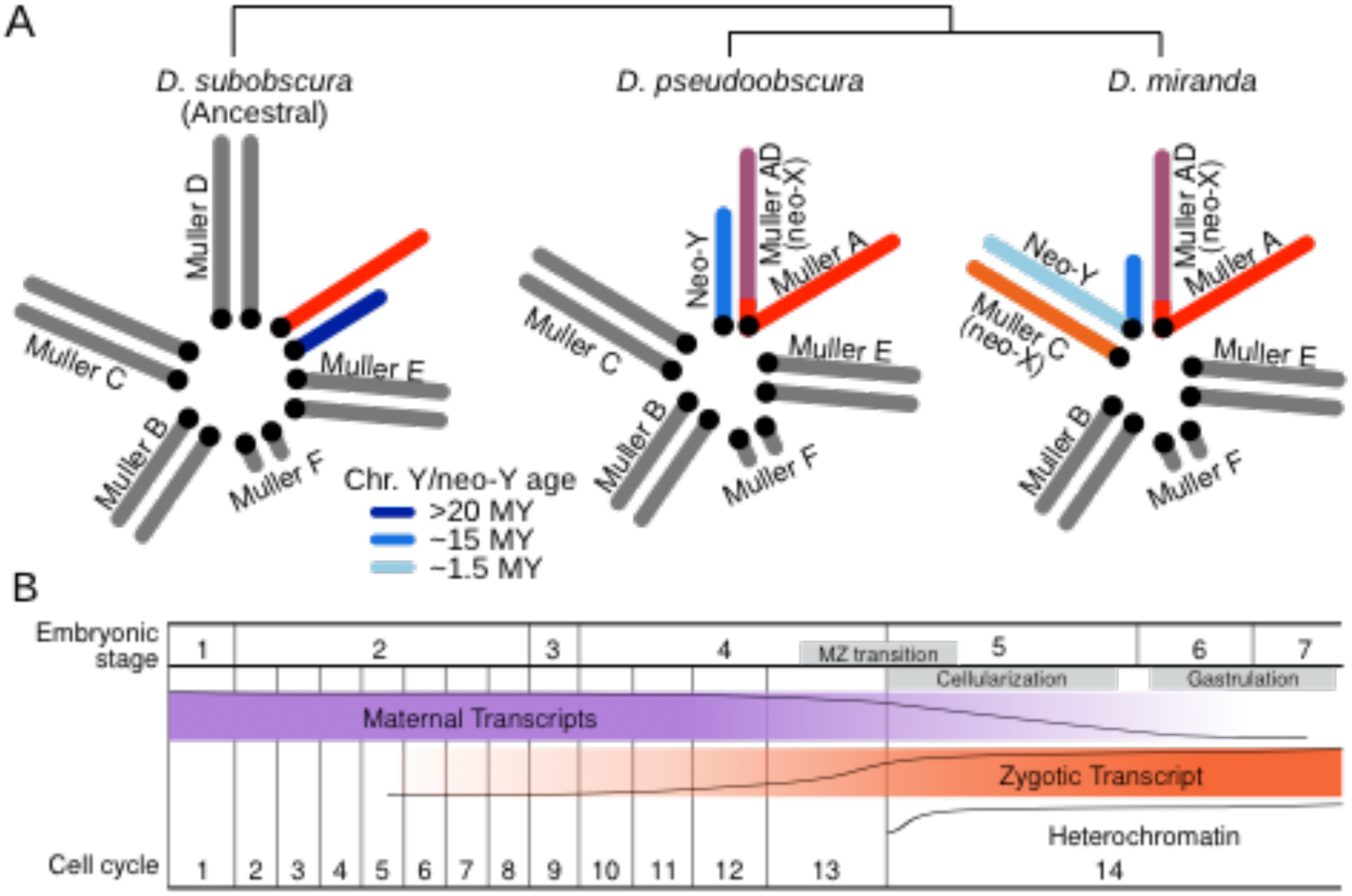
Neo-Y chromosome emergence and transcriptional activity during early embryogenesis in *Drosophila*. **A**. The male karyotypes of three species in the obscura species group showing three different Y and neo-Y (blue) chromosomes configurations. Chromosome arms are labeled as Muller Elements. The X/neo-X chromosome are in warm colors and autosomes in gray. *D. subobscura* represents the ancestral karyotype. *D. pseudoobscura* has a highly degenerate and old neo-Y that formed 15 MYA after the fusion of the X (Muller A) with Muller D, creating a neo-X chromosome. *D. miranda* has a young neo-Y that formed after Muller-C fused with the Y ∼1.5 MYA. The unfused Muller C then became the neo-X. **B**. Schematic representation of transcriptional changes and heterochromatin regulation in early embryonic development. The approximate abundance of maternal transcripts (purple), zygotic expression (orange), and repressive heterochromatin (blue) are depicted.

## Results and Discussion

### Elevated zygotic TE transcripts in male embryos

To study the dynamics of early repeat activation in *D. pseudoobscura* and *D. miranda*, we combine gene expression and chromatin profiles in single embryos during early development. In Drosophila, zygotic transcription begins at the pre-blastoderm (stage 2) and gradually increases; by the end of stage 4 (syncytial blastoderm), widespread zygotic transcription is observed (the maternal-to-zygotic (MZ) transition, see **Fig. 1B**) (Pritchard and Schubiger 1996; Lécuyer *et al*. 2007; Lott *et al*. 2014; Yuan and O’Farrell 2016; Strom *et al*. 2017). We analyzed transcriptomes of replicate single sexed embryos of *D. pseudoobscura* and *D. miranda* that have been developmentally staged from embryonic stage 2 through 12 (**Fig. 2A**; **Fig. S1; Fig. S2**; **Table S1**, (Lott *et al*. 2014)). Hierarchical clustering of the samples by their TE transcript abundance divides the embryos into two distinct groups that coincide with the MZ transition (**Fig. 2B**). Prior to the onset of widespread zygotic transcription (stages 2 and 4), TE transcript profiles are highly correlated between stages of the same species (**Fig. 2B**). As expected, female and male embryos are highly similar as the transcript pool is predominated by maternally deposited RNAs. After the MZ transition, samples form clades separated by species and sex (**Fig. 2B**). Interestingly, while *D. pseudoobscura* female and male samples are highly correlated and clustered, TE transcription profiles in *D. miranda* are less similar between sexes, and females of *D. miranda* group more closely to *D. pseudoobscura* (**Fig. 2B**).

**Figure 2.**
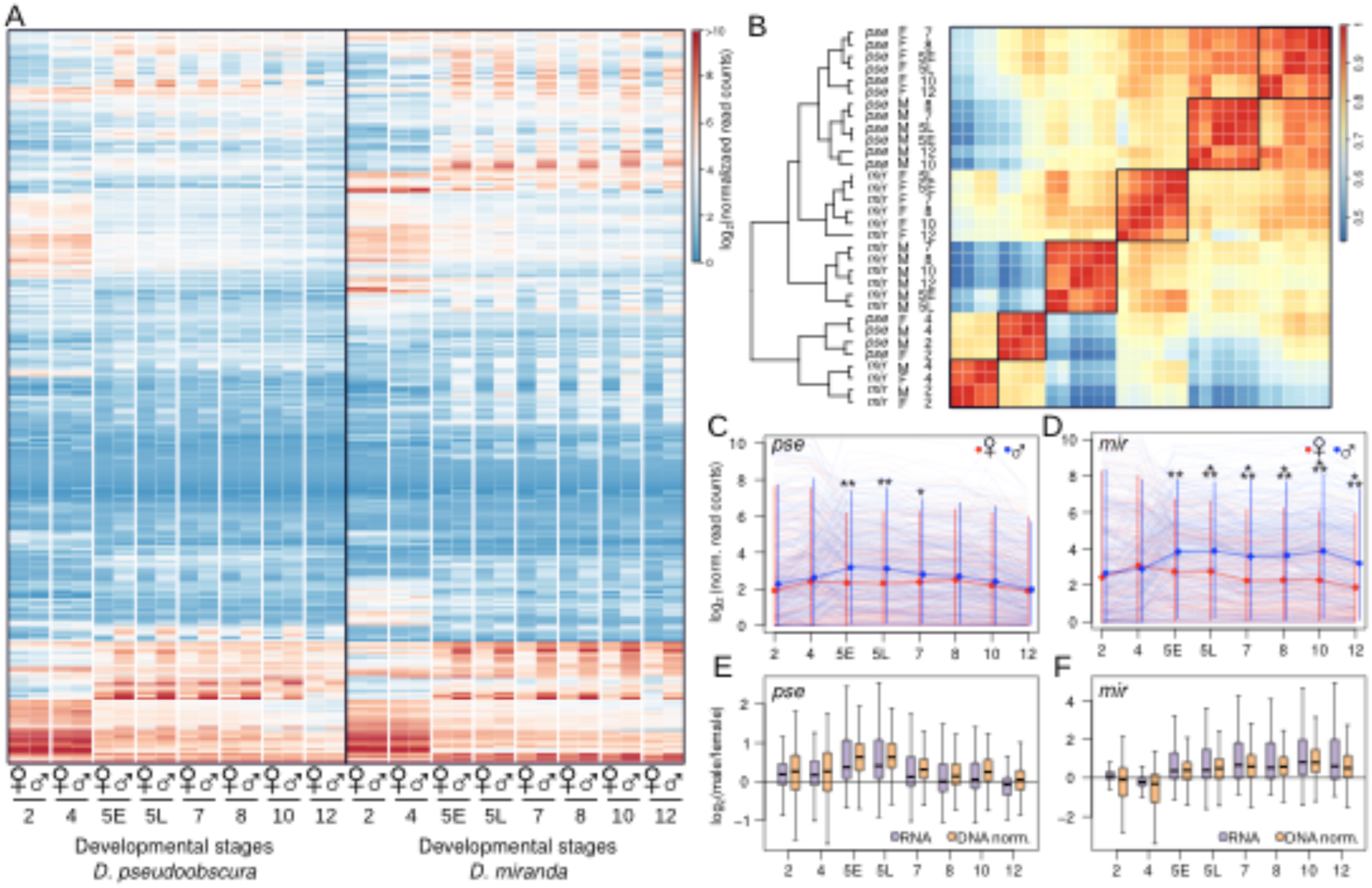
Sex-specific transcriptional regime of transposable elements across early embryogenesis. **A**. Heatmap showing the normalized read counts mapping to TEs from developmentally staged and sexed embryos (average of 3 embryos) from *D. pseudoobscura* and *D. miranda*. The list of TE names can be found in **Fig. S1**. For heatmap in FPKM, see **Fig. S2. B**. Heatmap depicting the correlation of TE transcript abundance between species, sex and developmental stages. Samples were ordered by hierarchical clustering. **C,D**. TE transcript abundance across developmental stages and sexes in *D. pseudoobscura* and *D. miranda*. Light lines depict TE transcript abundance for individual TEs, and dark lines depict the median with 95% confidence intervals demarcated by vertical lines. *, **, and *** denote significant differences between males and females with p-values of <0.01, <0.0001, and <0.000001, respectively (Wilcoxon’s Rank Sum Test). **E,F**. Distribution of the fold-differences in TE transcript abundance (purple) between sexes in log_2_ scale. RNA-seq read counts at TEs are further normalized by their DNA-seq read counts to account for copy number, and the fold-differences between sexes are depicted.

The sex differences after the MZ transition appear to be driven in part by higher TE expression in males in both species (**Fig. 2A,C,D; Fig. S3**); no such differences are seen at autosomal genes (**Fig. S4**). As zygotic expression increases, TEs are activated more highly in males than in females (**Fig. 2A**). *D. pseudoobscura* males have significantly higher TE expression than females immediately after the transition; at late stage 5 where the difference is greatest, males have on average 1.68-fold higher TE expression than females (**Fig. 2C**). TE expression in males then gradually decreases throughout development resulting in similar expression levels between the sexes at stage 12 (**Fig, 2A,C,E**). This suggests that efficient silencing mechanisms are established by then, in both sexes (see below). In *D. miranda*, TE transcripts are generally more abundant than in *D. pseudoobscura* (**Fig. 2A,C,D**) and males similarly show significantly higher TE expression than females. However, elevated TE expression is maintained in *D. miranda* males throughout early development (**Fig. 2A,D**). At stages 10 and 12, *D. miranda* males have on average more than twice the TE expression of females (2.37 and 2.04-fold, respectively; **Fig. 2D,F**), suggesting that TEs may be evading silencing in *D. miranda* males. Notably, elevated transcript abundance in males is not simply due to the higher copy number of repeats; after normalizing the TE read counts with their copy numbers in males and females, higher transcript abundance in males persists in both species (**Fig. 2E,F**).

### Y-linked TEs drive elevated TE expression in males

The presence of the Y and neo-Y chromosomes in males substantially increases the repeat content of the cell (Brown *et al*. 2020a). Elevated TE expression in males can either result from misregulation of Y-linked TEs or global reduction in repeat suppression (Brown *et al*. 2020a). Given the repetitive nature of TEs, it is typically not possible to pinpoint the specific genomic copy from which a TE transcript originates, especially for highly active families with large numbers of identical insertions. Instead, we identified TE families preferentially located on the Y chromosome based on their relative abundance in males vs. females (2-fold or higher mapping of DNA-seq reads); this resulted in 20 and 79 Y-enriched TEs in *D. pseudoobscura* and *D. miranda*, respectively (out of a total of 303 TEs; **Fig. 3A,B**). In both species, Y-enriched TEs are significantly more highly expressed in males after the MZ transition compared to females, and the magnitude of male-bias is also significantly higher than for the remaining TEs (**Fig. 3C-F**). These results are consistent with misregulation of Y-linked TEs driving elevated repeat expression in males. TEs not classified as Y-biased nonetheless show moderately male-biased expression, suggesting that global TE regulation may also be affected in males (**Fig. 3E-F**). The extent of male biased expression of Y-enriched TEs decreases through development in *D. pseudoobscura* (**Fig. 3G**) as their transcript abundance declines in both females and males (**Fig. 3C,E,G**). Only a small set of Y-enriched TEs do not drop to the same levels as in females, suggesting that some Y-linked copies may not be fully silenced (**Fig. S3**). In contrast, a large number of Y-enriched TEs maintain their high expression level throughout development in male *D. miranda* (**Fig. 3D; Fig. S3**); in fact, they become more male biased with time (**Fig. 3H**), largely due to decreased TE expression in females (**Fig. 3D,F**). Thus, Y-linked TEs appear to be poorly suppressed in *D. miranda* causing persistently elevated expression in males.

**Figure 3.**
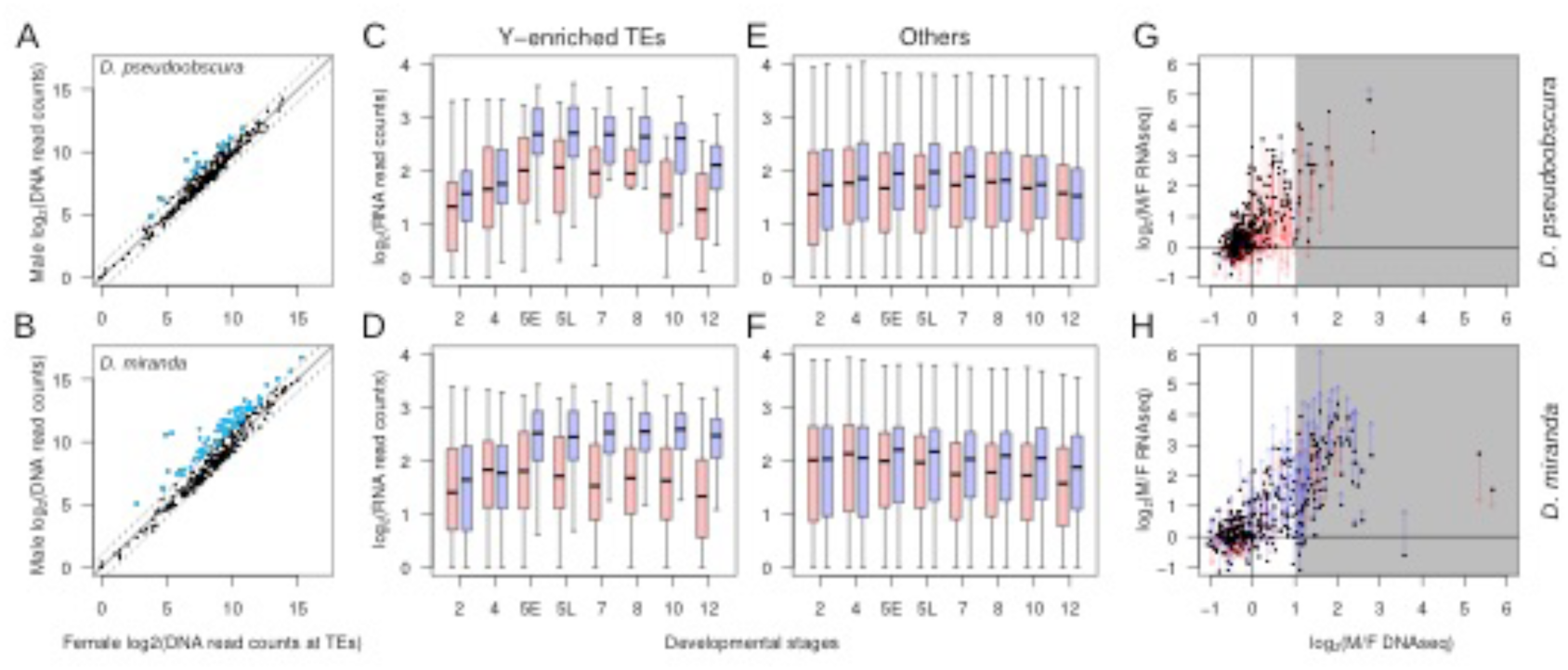
Y-linked TEs drive male-biased expression in both *D. pseudoobscura* and *D. miranda*. **A,B**. Regression between female and male DNA-seq read counts mapping to TEs. TEs with more than twice the abundance in males are deemed Y-enriched (blue). Solid diagonal delineate the identity line, dotted lines delineate ±2-fold differences. **C-F** Boxplots depict the transcript abundances of either the Y-enriched (**C, D**) or remaining (**E, F**) TEs between the sexes (female in red, male in blue) and across developmental stages. **G,H**. Fold-difference between female and male DNA-seq read counts plotted against that from RNA-seq read counts at stage 5. Gray background demarcate cutoff for Y-enriched TEs. Fold-expression difference between sexes for each TE are shown at developmental stage 5 (dots) and stage 12 (arrowheads). The color of the vertical line denotes whether the expression of a TE becomes less (red) or more (blue) male-biased across development.

### Reduced levels of heterochromatin at TEs on the *D. miranda* neo-Y

To determine if incomplete epigenetic silencing may drive elevated expression of Y-linked TEs after the MZ transition, we characterized genome-wide enrichment profiles of the repressive chromatin mark H3K9me3 in single sexed embryos at stages 5 and 7 (**Fig. 4A,B**). These two time points reflect the transition of initiation of heterochromatin (at the onset of cellularization of the blastoderm in early stage 5) towards maturation into a stable, repressive chromatin compartment (gastrulation of the embryo at stage 7) (Vlassova *et al*. 1991; Lu *et al*. 1998; Yuan and O’Farrell 2016). Overall, H3K9me3 is enriched at repeat-rich regions in both species at both stages, including the Y/neo-Y chromosomes and the pericentromeres (**Fig. 4A,B**). As expected, H3K9me3 enrichment is higher at stage 7, consistent with the progressive establishment and spreading of heterochromatin after the MZ transition (Vlassova *et al*. 1991; Lu *et al*. 1998; Yuan and O’Farrell 2016). Therefore, this suggests that decreasing male-biased TE expression across development in *D. pseudoobscura* likely results from increased heterochromatic suppression, and efficient silencing of Y-linked TEs (**Fig. 4A**). In contrast, the neo-Y chromosome of *D. miranda* does not appear to become fully heterochromatinized despite containing tens of thousands of TEs (**Fig. 4B**). Chromosome-wide H3K9me3 enrichment levels on the neo-Y are markedly less than that of pericentric heterochromatin at the X and autosomes, at both developmental stages, even though their repeat content is similar (**Fig. 4B, Fig. S5**).

**Figure 4.**
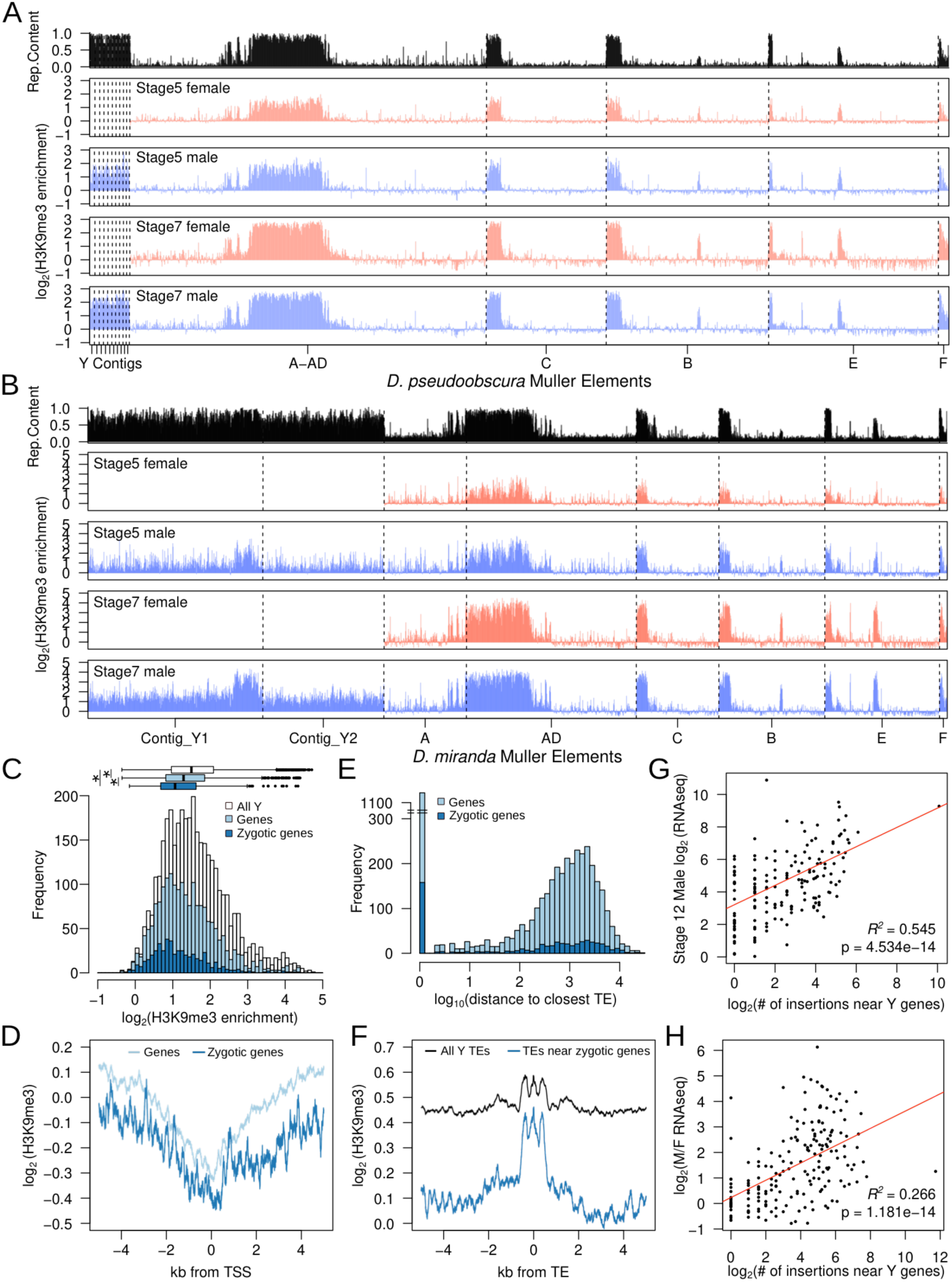
The heterochromatin landscape on the neo-Y. **A, B**. *D. pseudoobscura* and *D. miranda* H3K9me3 enrichment between sexes and across stages 5 and 7. The repeat content of the chromosomes are depicted at the top (in black). Chromosomes and contigs are separated by vertical dotted lines. For H3K9me3 enrichment estimation using only uniquely mapped reads, see **Fig. S5. C**. Distribution of H3K9me3 enrichment in 50kb windows on the neo-Y. Windows overlapping annotated genes and zygotically expressed genes are in light and dark blue, respectively. **D**. Median H3K9me3 enrichment ±5Kb around the transcription start site of all (light blue) and zygotically expressed (dark blue) genes on the neo-Y. **E**. Distance between genes and the nearest TE on the neo-Y. 0 indicates the TE is within a gene. **F**. Median H3K9me3 enrichment around all annotated TEs and those within ±5Kb of a gene on the neo-Y. **G, H**. The number of times TEs are found to have inserted near zygotically expressed genes on the neo-Y (within ±5Kb) is plotted against the TE transcript abundance (**G**) and the fold difference of TE transcript abundance between males and females (**H**) at stage 12. Line of best fit is in red; Pearson’s *R* of the regressions and the significance of the correlations are labeled. Only TEs found on the neo-Y chromosome are plotted (n=198).

### Transcription of neo-Y genes impedes heterochromatin formation at TEs

The genome architecture of the neo-Y differs from pericentromeres; despite consisting mostly of repetitive DNA, the neo-Y still contains thousands of functional genes with 6448 genes annotated in the current assembly (Mahajan *et al*. 2018; Bachtrog *et al*. 2019). Active transcription at neo-Y genes may impede heterochromatin formation, creating islands of euchromatin across the neo-Y (Allshire and Madhani 2018). Indeed, while heterochromatic Y chromosomes do not form polytene chromosomes in Drosophila (Leach *et al*. 2000), segments of the neo-Y chromosome maintain euchromatin-like banding patterns in polytene spreads interspersed with under-replicated heterochromatin (Macknight and COOPER 1944). TEs may therefore exploit these euchromatic environments to maintain elevated activities. Supporting this hypothesis, segments of the neo-Y adjacent to active genes are less heterochromatic: Neo-Y windows overlapping annotated genes have significantly lower H3K9me3 enrichment compared to all neo-Y windows (Wilcoxon rank sum test p < 2.2e-16; **Fig. 4C**), and windows with zygotically expressed Y-linked genes have even less H3K9me3 enrichment (Wilcoxon rank sum test p = 4.689e-07; **Fig. 4C**). H3K9me3 levels are depleted near the transcription start sites of both Y-linked and zygotically expressed Y-linked genes and gradually increase distally (**Fig. 4D**).

To determine whether TEs profit from reduced silencing near Y-linked genes, we evaluated the distribution of genes and TE insertions on the neo-Y. Y-linked genes are on average only 512bp away from the closest TEs and 25.4% have at least one insertion within the gene (and 27.6% of zygotically expressed Y genes), presumably in the introns and UTRs (**Fig. 4E**). In contrast, genes and TEs are significantly farther apart on autosomes, with an average distance of 4127bp and 7.7% of autosomal genes have internal TE insertions (Wilcoxon rank sum test p < 2.2e-16; **Fig. S6**). Despite elevated H3K9me3 enrichment, TEs near Y-linked zygotically expressed genes are less enriched than TEs across the entire chromosome (**Fig. 4F**). Consistently, TEs are expressed more highly in males if they have a larger number of insertions around (±5Kb) zygotically expressed neo-Y genes (p = 4.53e-14; **Fig. 4G**), and these TEs show higher male-bias in their expression (p = 4.87e-05; **Fig. 4H**). Therefore, TEs neighboring transcribed neo-Y linked genes are poorly suppressed and likely a main source of the persistently elevated expression in *D. miranda* males. Notably, while TEs near zygotically expressed genes show reduced H3K9me3 levels, they are nevertheless enriched for heterochromatin, suggesting some extent of epigenetic silencing of TEs. The deposition and spreading of silencing heterochromatin at TEs near euchromatic genes is deleterious and euchromatic TE insertions are under purifying selection in *Drosophila* (Lee 2015). The close proximity of thousands of genes and TEs on the neo-Y therefore results in an epigenetic conflict between silencing TEs via heterochromatin formation while maintaining the activity of functionally important genes during development.

### Increased rates of TE insertions in *D. miranda* males

Elevated TE expression could lead to increased rates of TE insertions. To test if differences in TE activity result in sex-specific differences in TE movement, we identified novel TE insertions in male and female embryos by deep sequencing, leveraging the input DNA reads from our single embryo ChIP-seq data. Insertions were defined by paired-end reads where one read mapped uniquely to the genome and the other to a TE (Jiang *et al*. 2015; Treiber and Waddell 2017)(**Fig 5A**). To avoid capturing chimeric reads, we required that both the 5’ and 3’ junctions of insertions were identified (**Fig 5A,B**), and TEs found in more than one sample are discarded to ensure only novel insertions are counted. As any *de novo* insertion is likely found in a tiny fraction of cells, our approach is a highly conservative estimate of the number of total insertion events (**Table S3**). To ensure that our approach is robust to artifacts, we tested it using exon sequences instead of repeats; indeed, no more than 4 “insertions” are called across any library (and a median of 0 “insertions” per library; **Table S4**).

**Figure 5.**
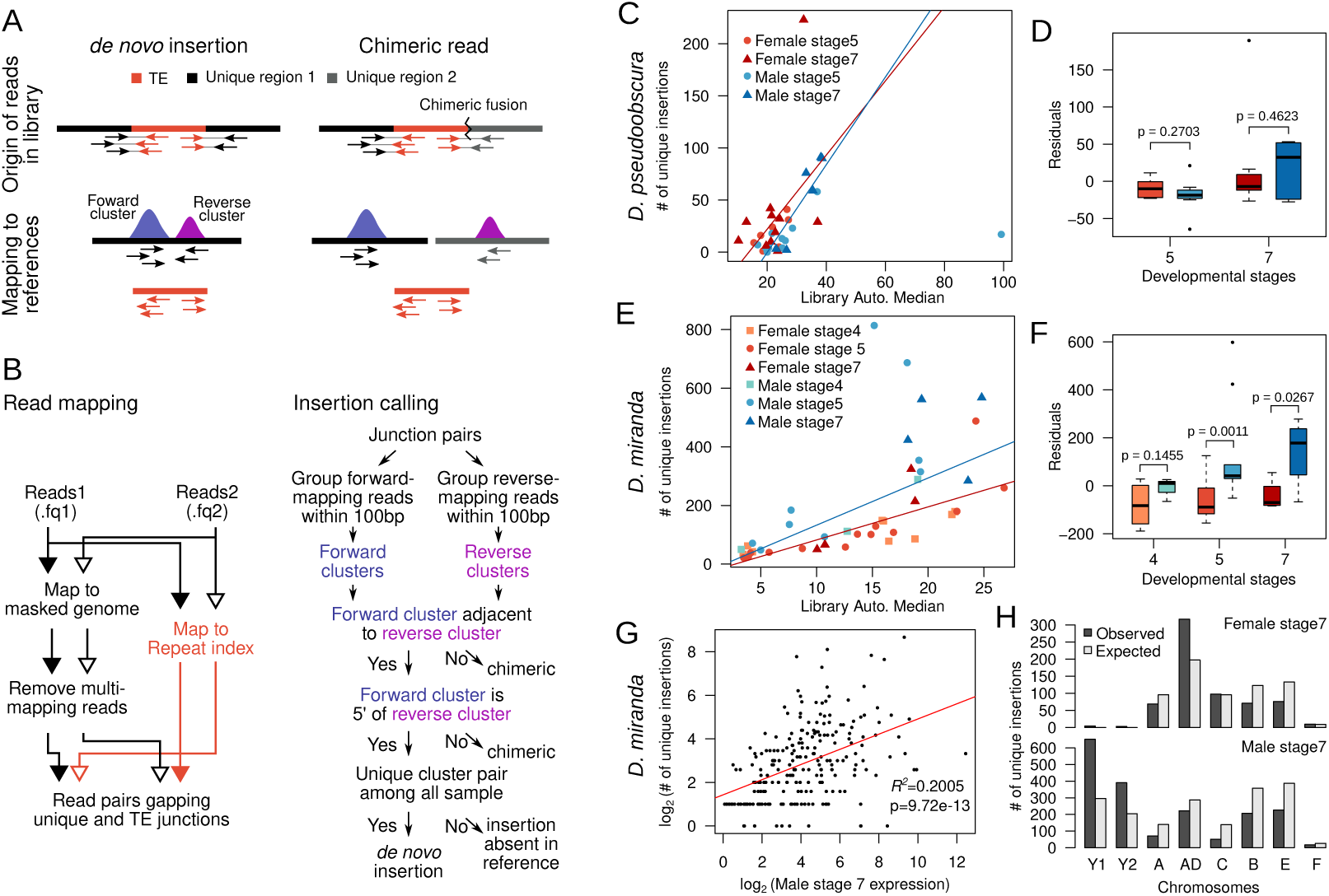
*D. miranda* males show more *de novo* TE insertions than females. **A**. Schematic diagram of rationale behind TE insertion identification. **B**. Computational pipeline implementing the strategy in A. The left panel outlines the mapping strategy in identify mate pairs at insertion junctions. The right panel outlines the filtering strategy to identify *de novo* TE insertions. See Table S3 for summary of insertions filtered and identified for each library. **C**. The numbers of *de novo* TE insertions identified in *D. pseudooboscura* embryos are plotted against the autosomal median coverages. Lines of best fit are shown for the male (blues) and female (reds) samples, separately. **D**. The effect of library size is removed by regression of TE insertion numbers against library coverage and taking the residuals. Boxplot depicts the residuals of the regression binned by sex and developmental stages. P-values are labeled above the comparisons (Wilcoxon’s Rank Sum Test). **E,F**. Same as C and D but for *D. miranda* embryos. **G**. The number of *de novo* insertions across all male *D. miranda* stage 7 embryos is plotted against the transcript abundance. Line of best fit is in red and *R*^*2*^ of the regression and the significance of the correlation are shown. Only TEs with de novo insertions are plotted. **H**. The observed and expected distribution of insertions across the chromosomes are depicted in females (top) and males (bottom).

We identified a total of 1054 and 8191 insertions across 37 *D. pseudoobscura* and 42 *D. miranda* single embryo libraries (**Table S3**). The majority of these insertions are likely somatic as the number of somatic cells are at least three orders of magnitude more numerous than the pole (germ) cells at these embryonic stages. As expected, the number of TE insertions identified from the embryos is strongly dependent on the sequencing depth (**Fig. 5C**,E). For each of the two species, we fitted an ANOVA model with median coverage, developmental stage, and sex as independent variables, and number of insertions as the response variable. While library coverage has the strongest effect in both species, we find a significant effect of sex in *D. miranda*, but not in *D. pseudoobscura* (**Table S5**). Additionally, developmental stage also shows a strong effect in both species reflecting the increase in TE activities through the MZ transition and the increasing number of cells where insertions can occur. After removing the effect of library coverage, we observed significantly more TE insertions in males than in females for stage 5 and 7 embryos in *D. miranda* but not *D. pseudoobscura* (**Fig. 5D,F, Fig. S7**). Increased rates of TE insertions in *D. miranda* males are also observed when considering only autosomal insertions (**Fig. S8**). The magnitude of difference is greater in stage 7 embryos, likely due to the increased amount of time and cells in which *de novo* mutations can occur (**Fig. 5F**). Female and male embryos at stage 4 have similarly low numbers of insertions (**Fig. 5F**), consistent with the absence of male-biased TE expression prior to the MZ transition. As expected, *de novo* TE insertions resulted from repeats that show higher expression in early embryos (Charlesworth and Langley 1986), and the number of insertions summed across all embryos of the same sex and stage is significantly correlated with their transcript abundance (**Fig. 5G**). Altogether, these results reveal that elevated TE expression in *D. miranda* males is associated with higher rates of TE insertions in males compared to females. Interestingly, we find that insertions are non-randomly distributed across chromosomes. In both males and females, there are significantly fewer autosomal insertions than expected based on the chromosomal sizes (Fisher’s Exact Test, p < 0.0001; Figure 5F). TE insertions are significantly overrepresented on Muller-AD in females and the neo-Y in males (FET, p < 0.0001 for both; **Fig. 5F**), and either transposition bias or selection could contribute to the non-random spatial distribution of TEs (Bousios *et al*. 2020). The elevated rate of somatic TE insertions in *D. miranda* males is expected to impose a deleterious fitness cost uniquely to males, and may contribute to the female-biased sex ratio found in this species (Dobzhansky 1935), and shorter lifespan of *D. miranda* males compared to females (Nguyen and Bachtrog 2020). Additionally, if insertions rates are also higher in the male germline, the species as a whole is expected to have a higher mutational TE burden. Concordantly, the *D. miranda* genome overall has higher repeat content than its close relative *D. pseudoobscura*, even outside of its neo-sex chromosomes (Bracewell *et al*. 2019).

### Model for neo-Y toxicity and TE accumulation

Y chromosomes have evolved repeatedly in different species from a pair of ordinary autosomes. Y evolution is typically characterized by progressive gene loss, an accumulation of repetitive DNA, and heterochromatin formation (Bachtrog 2013). Here, we show that nascent Y chromosomes can form a “genomic liability” for males, especially if epigenomic defense mechanisms are compromised. This occurs in early development, where the zygotic genome is reprogrammed to create a set of totipotent cells capable of generating a new organism. Heterochromatin loss also occurs in old individuals, and Y chromosomes can contribute to faster male aging in Drosophila (Brown *et al*. 2020b). We show that incomplete silencing of Y-linked TEs in early development results in a surge of repeat expression in males, resulting in more somatic TE insertions. Repeat-rich Y chromosomes that still contain functional genes create a dilemma, as actively transcribed euchromatin antagonizes heterochromatin assembly (Allshire and Madhani 2018). Thus, competition between the opposing mechanisms of heterochromatin formation and transcription likely explains the incomplete silencing of TEs on the transcriptionally active neo-Y in *D. miranda*. While the accumulation of repetitive elements on the Y chromosome appears to be near universal during sex chromosome evolution, our study reveals an unappreciated aspect of this process (**Fig. 6**). The conflict between genic expression and TE silencing on a nascent Y creates a ‘toxic environment’ elevating the mutational burden in the male genome. This discord is maximized on Y chromosomes of intermediate age that still contain an appreciable number of genes but also a high repeat density. Resolution of this conflict may select for the adaptive degeneration of remaining protein-coding genes on the Y and further repeat accumulation to strengthen epigenetic silencing, thereby reducing the toxicity of the Y. Epigenetic conflicts therefore represent a novel mechanism driving the degeneration of the Y chromosome.

**Figure 6.**
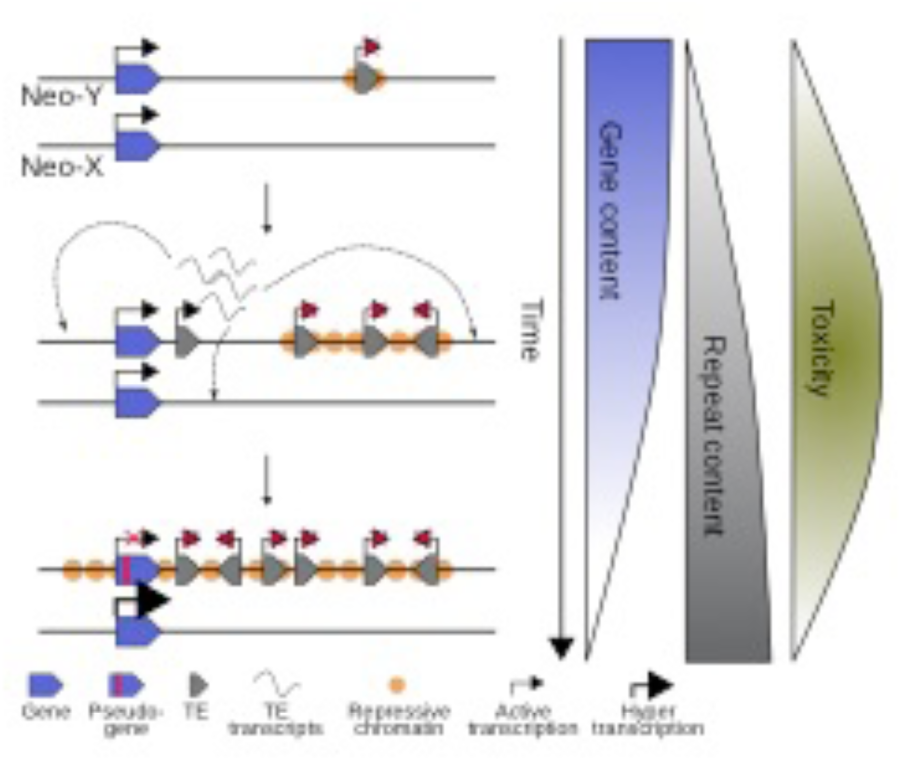
Toxic Y chromosome and adaptive Y degeneration. Epigenetic conflicts between host-specific genes (blue) and selfish genetic elements (grey) are maximized on Y chromosomes of intermediate age (maximum toxicity). Active transcription of Y-linked genes impedes heterochromatin formation at repeats that accumulate on degenerating Y chromosomes and their mobilization can create a mutational burden in males (middle). Selective pressure to efficiently silence a toxic Y chromosome and dosage compensation of the X homologs could drive adaptive degeneration of Y genes.

## Methods

### Fly strains

We used *D. pseudoobscura* strain SS-R2 and *D. miranda* strain MSH22 kept at 18°C (the preferred temperature for these species and the same temperature they were reared when the RNA-seq data and ChIP-seq data was generated) and *D. melanogaster* strain Oregon-R kept at 25°C.

### RNA-seq data analysis

We used published RNA-seq data (Lott *et al*. 2014) to analyze sex-specific repeat expression during embryogenesis. For read counts at genes, pair-end reads were mapped to the *D. pseudoobscura* (Bracewell *et al*. 2019)or *D. miranda* (Mahajan *et al*. 2018) genomes using bwa mem (v0.7.15) (Li and Durbin 2009) on default settings and sorted with samtools (v1.5) (Li *et al*. 2009). We then used featureCounts (v1.6.2) from the Subread package (Liao *et al*. 2014) to determine the number of reads mapping to annotated genes of the two genomes. For counts at TEs, we mapped reads with bwa mem to the TE library specific to the *pseudoobscura* species subgroup generated by (Hill and Betancourt 2018). Reads mapping to each repeat entry were then tallied from the sam files. We also used bowtie2 (v2.3.4.1), which generated similar patterns in sex-biased expression indicating that our results are robust to mapping strategies (**Fig. S9**). We normalized the gene and TE read counts by the median read counts at autosomal genes to avoid the large contribution of sex-specific expression from the sex chromosome, especially the neo-Y. After normalization, one pseudocount is added to each gene. All hierarchical clustering and correlation procedures were conducted on the log_2_ transformed read counts. For sample information and mapping statistics, see Table S1.

### Chromatin immunoprecipitation and sequencing

We performed ChIP-seq using a protocol adapted from (Brind’Amour *et al*. 2015). *D. pseudoobscura* data were newly collected, and *D. miranda* data was downloaded from the SRA under BioProject PRJNA601450. Briefly, single embryos were homogenized with a pipette tip, and chromatin was digested for 7.5 minutes at 21°C using micrococcal nuclease (MNase) (New England Biolabs). We spiked DNA from *D. pseudoobscura* single embryos with DNA from *D. melanogaster* stage 7 (gastrulation) embryos so that each sample had 20% spike (*D. melanogaster*) DNA (i.e. one *D. melanogaster* embryo was used for four *D. pseudoobscura* embryos). We set aside 10% of each sample as input and incubated the remaining chromatin with Dynabeads Protein G (Invitrogen) for 2-6 hours. The H3K9me3 antibody (Diagenode, 1.65 ug/ul) was incubated for >3 hours with Dynabeads Protein G to bind the antibody to the beads, before adding it to the chromatin (0.25 ul per embryo) for overnight incubation. The chromatin-antibody-bead complexes were washed first with low-salt buffer, then with high-salt buffer. DNA was eluted from the chromatin-antibody-bead complexes by shaking at 65°C for 1-1.5 hours. We extracted DNA from our ChIP samples and from our input using a phenol/chloroform/isoamyl alcohol mixture, then cleaned the DNA with Agencourt AmpureXP beads before library preparation. Libraries were prepared using the ThruPLEX DNA-seq kit (Rubicon) followed by two rounds of cleaning with AmpureXP beads. 100bp paired-end sequencing of our samples was performed at the Vincent J. Coates Genomic Sequencing Laboratory at UC Berkeley.

### ChIP-seq data processing and normalization

ChIP-seq libraries for *D. miranda* were downloaded from the SRA under BioProject PRJNA601450. Like the *D. pseudoobscura* libraries they were spiked with D. melanogaster embryos. To distinguish the spike-in reads, pair-end reads were aligned using bwa mem to a concatenated reference combining the *D. melanogaster* genome (r6.12) with either the *D. pseudoobscura* or *D. miranda* genomes. Since bwa mem by default only reports the best alignment, spike-in reads are then extracted as those mapping to the D. melanogaster contigs. We used bedtool’s genomeCoverageBed (Quinlan and Hall 2010) to obtain per base coverage of the sample and spike-in reads after sorting with samtools sort. For library and mapping information see Table S2.

For the genome-wide coverage H3K9me3 enrichment in both spike-in and actual samples, enrichment at 50kb window is calculated as:

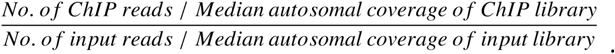

For spike-in normalization, we first generated an enrichment “reference” by averaging the enrichment of all spike-ins. Since all spike-ins should have the same enrichment, systematic differences between spike-ins are predominantly the result of different antibody pulldown efficiencies during library preparation. We used a quantile normalization procedure, matching the distribution of the enrichment values to that of the reference. The extent of change for each enrichment value after the normalization is then applied windows of the same enrichment value in the actual sample. In effect, if windows with enrichment of 1.5 in the spike-in is increased to 2 after quantile normalization, windows with enrichment of 1.5 in the actual sample will be increased to 2. For details on the procedure, see **Fig. S10**.

### H3K9me3 enrichment analysis

We used the R package IRanges (Lawrence *et al*. 2013) to infer overlaps and distances between windows with elevated H3K9me3, genes, and TEs. H3K9me3 enrichment around genes were inferred by extracting the per bp enrichment 5000bp up and down stream of the TSS of the relevant genes; the median enrichment at each position is plotted. For TEs, enrichments were extracted 5000bp up and down stream of the midpoint of the relevant TE annotation.

### TE annotation and repeat masking

We annotated the genomes of the two species and masked the repeats with the repeat library using RepeatMasker (Tempel 2012) (v3.3) with the following command:

~~~
RepeatMasker -norna -nolow -dir output.directory -gff -u -lib
TE.library.fasta genome.fa
~~~

### Identification of de novo TE insertions

Pair-end reads of inputs from the ChIP-seq experiments are mapped as single ends to their respective repeat-masked genomes and the repeat library. We first identified read pairs where one maps uniquely to the genome and the other maps only to the TE library, thus identifying pairs that flank insertion junctions. Reads of the same mapping orientation (forward or reverse strand mapping) that map <100bp of each other in the genome are deemed to be capturing the same junction. Because fusion of DNA fragments during library preparation can generate chimeric reads which can create such TE-to-unique junctions, we conservatively estimated insertions by further requiring that both the 5’ and 3’ junctions of the insertions must be identified. Thus, an insertion is only called if a forward mapping junction at the 5’ is followed by a reverse mapping junction <100bp away at the 3’. Note, since TEs can insert in either direction, we do not stipulate any directionality to reads mapping within the TEs. For true *de novo* insertions in the embryos, we removed all insertions that are found within < 50bp of any other insertion found across all samples.

With each of the two species, we used the following ANOVA model in R:

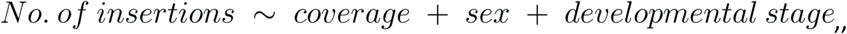

with coverage being the median autosomal coverage. For ANOVA summary statistics, see Table S5. To remove the effect of sequence depth, we used the linear regression model:

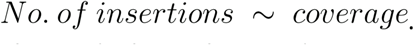

The residuals are then used to compare the difference in the amount of insertions due to sex and developmental stage. We calculated the expected number of TE insertions per chromosome by assuming uniform insertion rates; we multiplied the number of observed insertions genome-wide by the size of the chromosomes (in the genome assembly) proportional to the diploid genome of females and males.

## Data availability

All the sequencing data have been posted on GenBank under BioProject PRJNA625074. All processed data files have been posted on Dryad.

## Code availability

Scripts used for normalization and insertion calls can be found on KW’s github page https://github.com/weikevinhc/heterochromatin.

## Acknowledgments

This work was supported by NIH grants (nos. R01GM076007, R01GM101255 and R01AG057029) to DB.

