## Supplementary Material for "Epigenetic conflict on a degenerating Y chromosome increases mutational burden in Drosophila males"

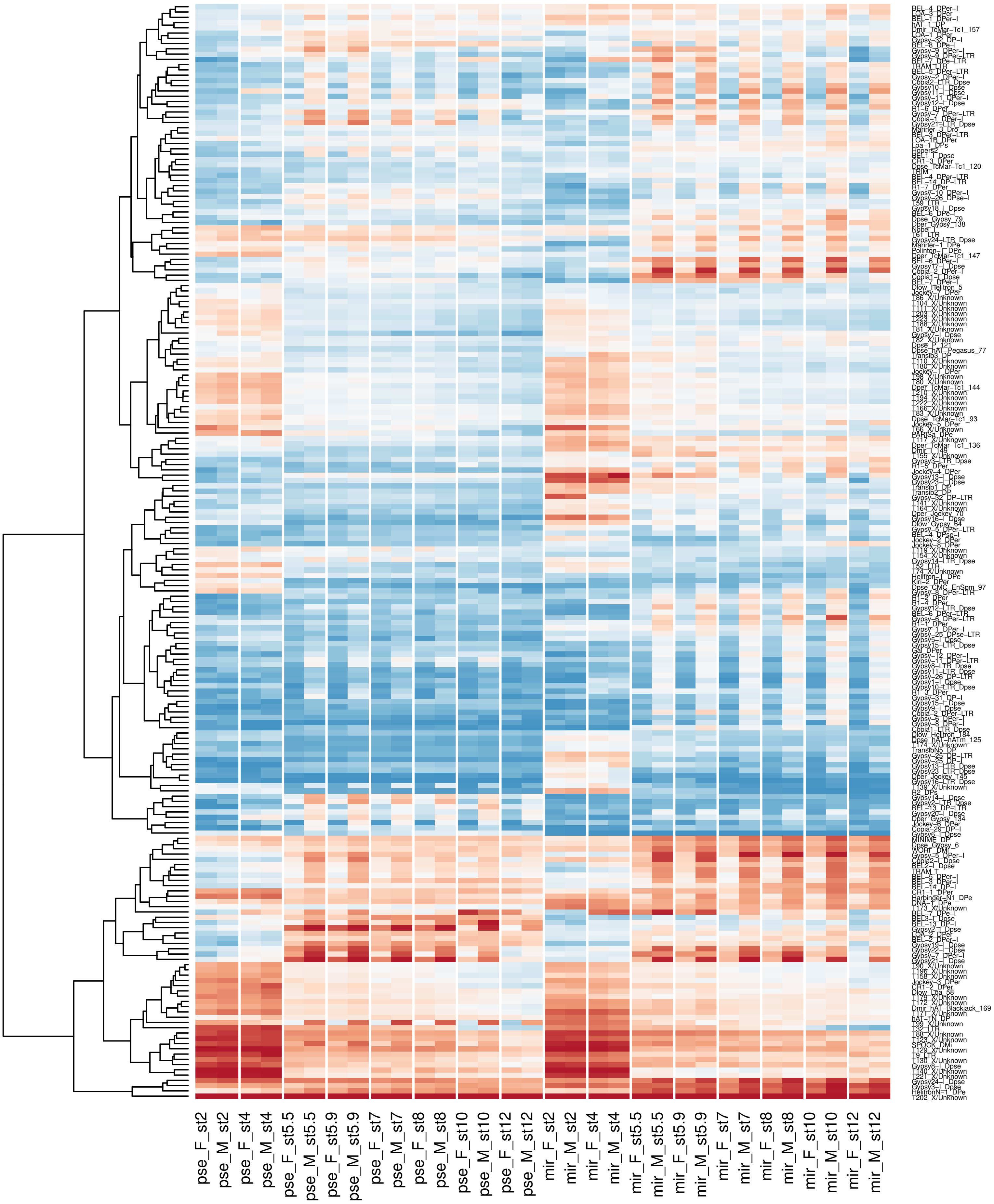


**Supplementary Figure S1.** Female and male TE expression across developmental stages in *D. pseudoobscura* and *D.miranda*. Normalized counts are averaged across replicates. TEs are labeled to thr right. TEs lowly expressed across all samples are removed.

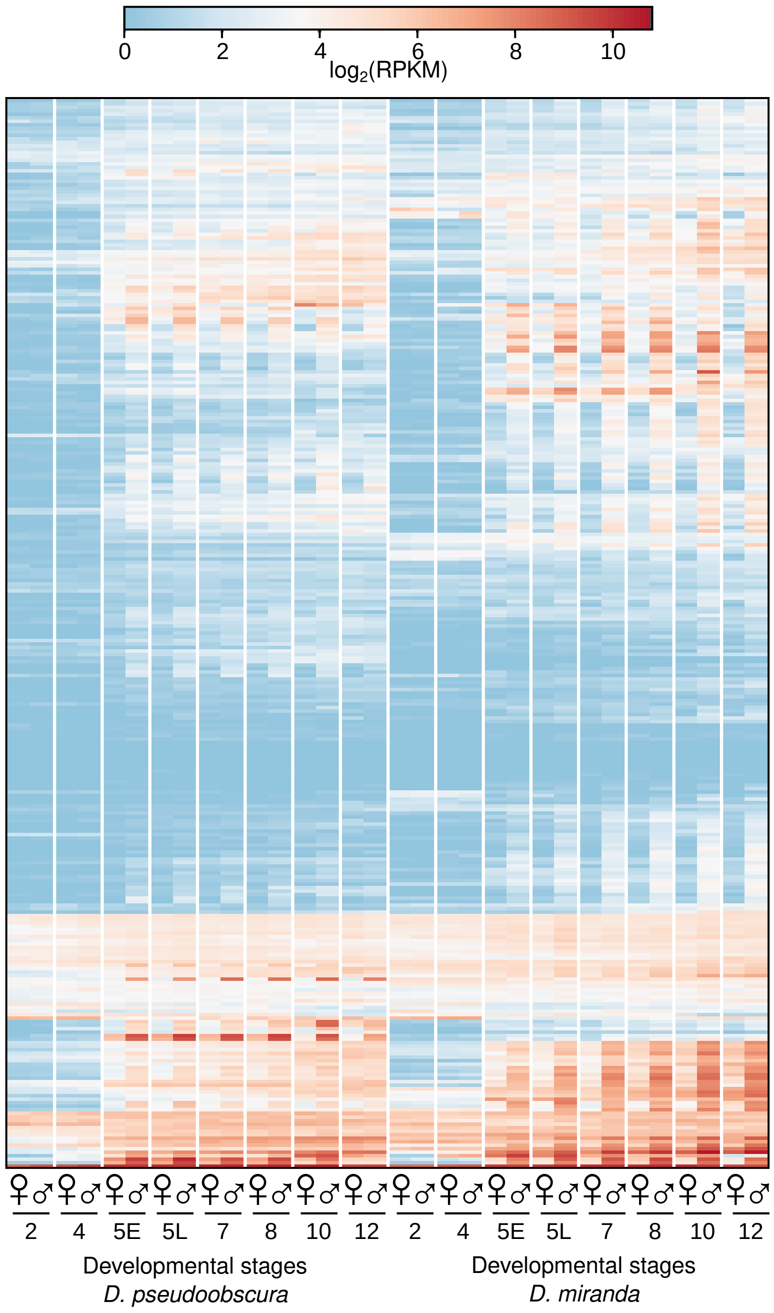


**Supplementary Figure 2. Transcript abundance with RPKM**. TE transcript abundance in RPKM across species, sex, and developmental stages.

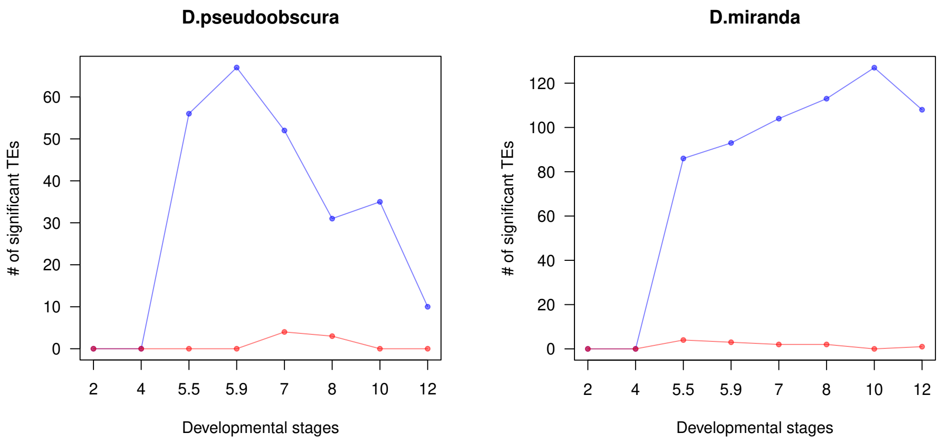


**Supplementary Figure 3.** Significantly differentially expressed TEs in *D. pseudoobscura* and *D. miranda*. The number of significantly male-biased (blue) and female-biased (red) TEs is plotted across the developmental stages. Significance is inferred using DESeq2 with FDR of 0.05.

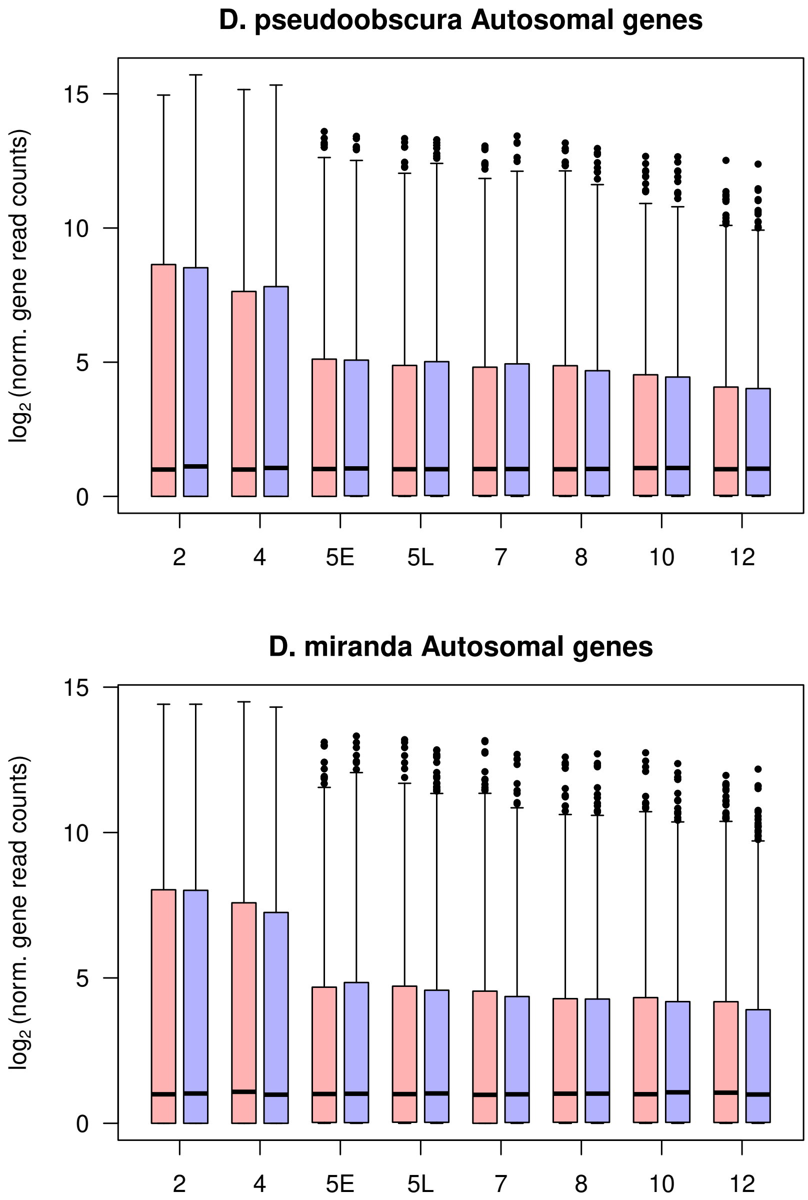


**Supplementary Figure S4.** Female (red) and male (male) autosomal gene expression across developmental stages in *D. pseudoobscura* (top) and *D.miranda* (bottom).

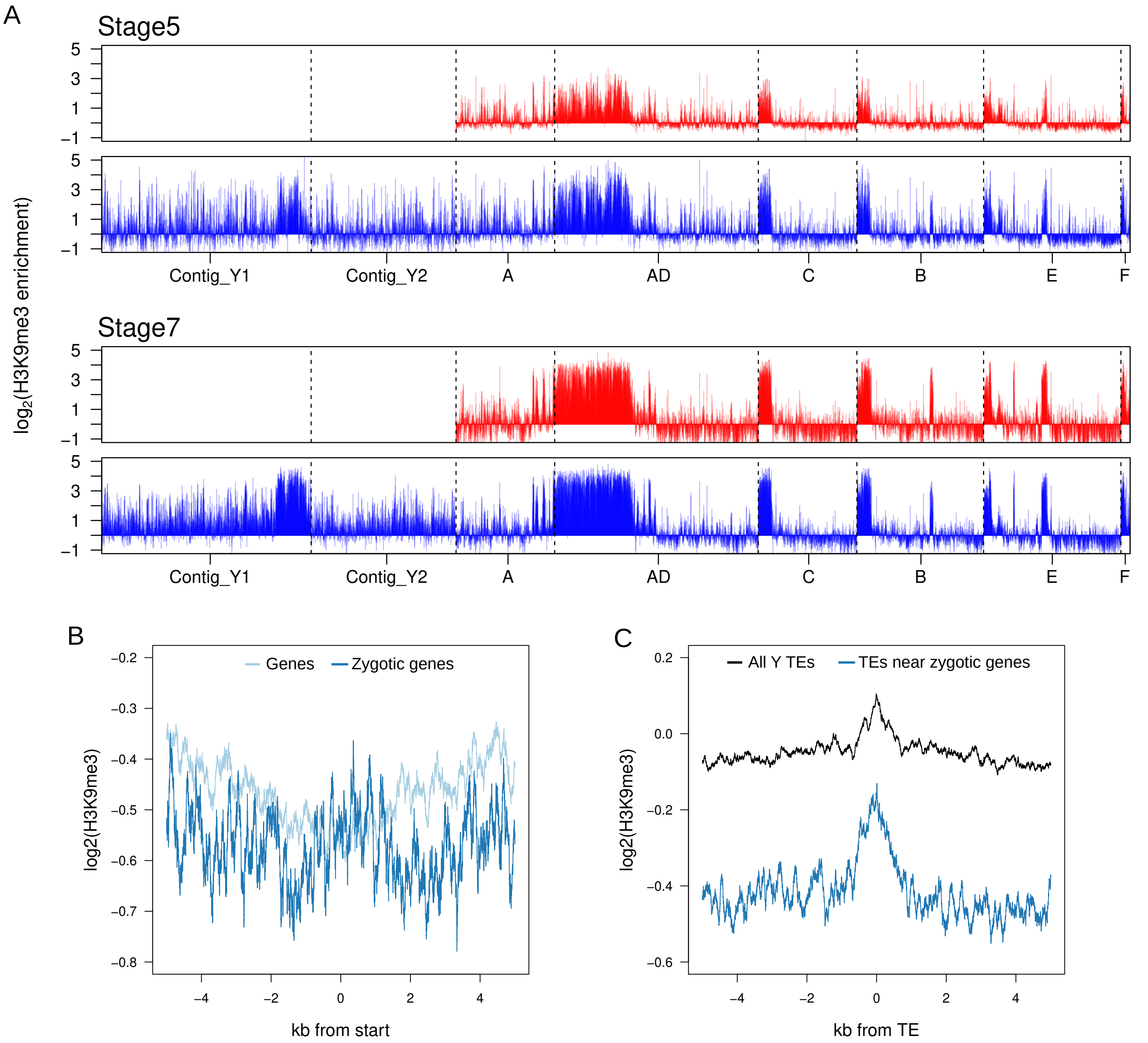


**Supplementary Figure 5. H3K9me3 enrichment in *D.miranda* using only uniquely mapping reads.** A. As with the enrichment profiles using non-uniquely mapping reads (Figure 4B), in males, the neo-Y chromsome has lower H3K9me3 enrichment when compared to the pericentromeric heterochromatin. B. Average enrichment around neo-Y genes. C. Average enrichment around neo-Y TEs.

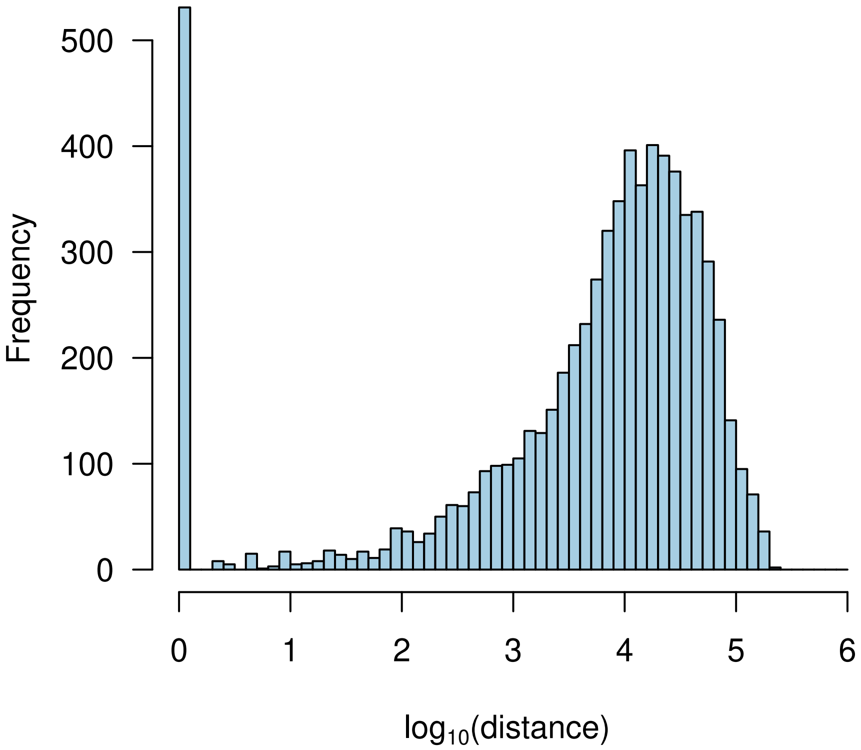


**Supplementary Figure S6.** Distribution of distance between autosomal genes and closest TEs. Average distance = 4127bp.

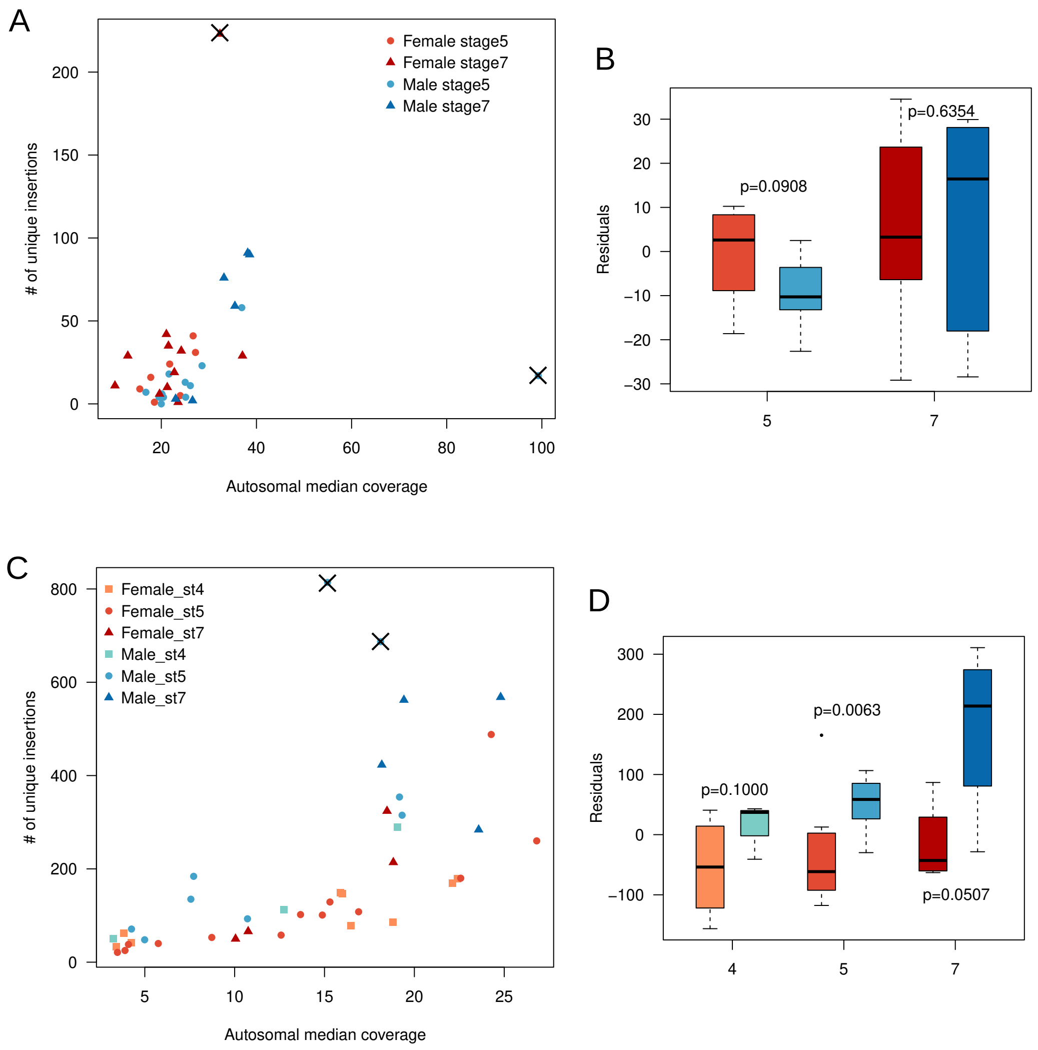


**Supplementary Figure 7. Insertions analyses after removal of outliers**. A. Number of unique insertions plotted against the median autosomal coverage of each D. pseudoobscura library, same as Figure 5C. Outliers are crossed out. B.To remove the effect of library size a linear regression acrossed the points in A; boxplots depict the residuals of the linear regression across different developmental stages and sex. Same as A and B, respectively, but for *D. miranda.*

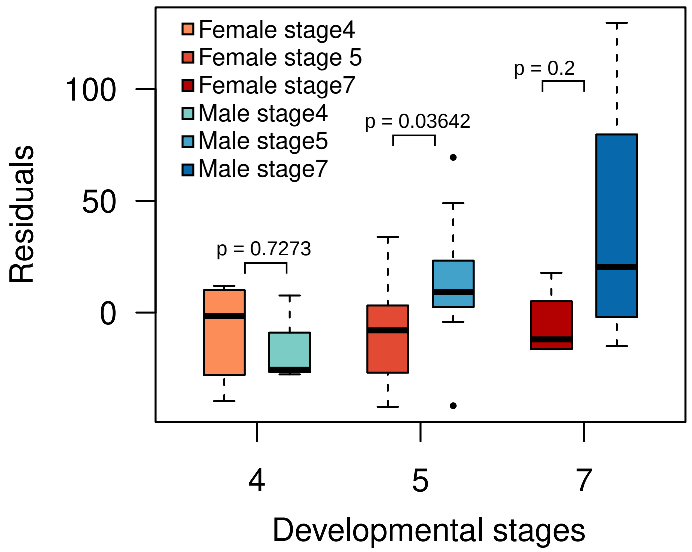


**Supplementary Figure S8.** de novo autosomal TE insertions in *D. miranda* across sex and age. Residuals are inferred from linear regression between library coverage and number of autosomal insertions. P-value inferred with Wilcoxon's Rank Sum test.

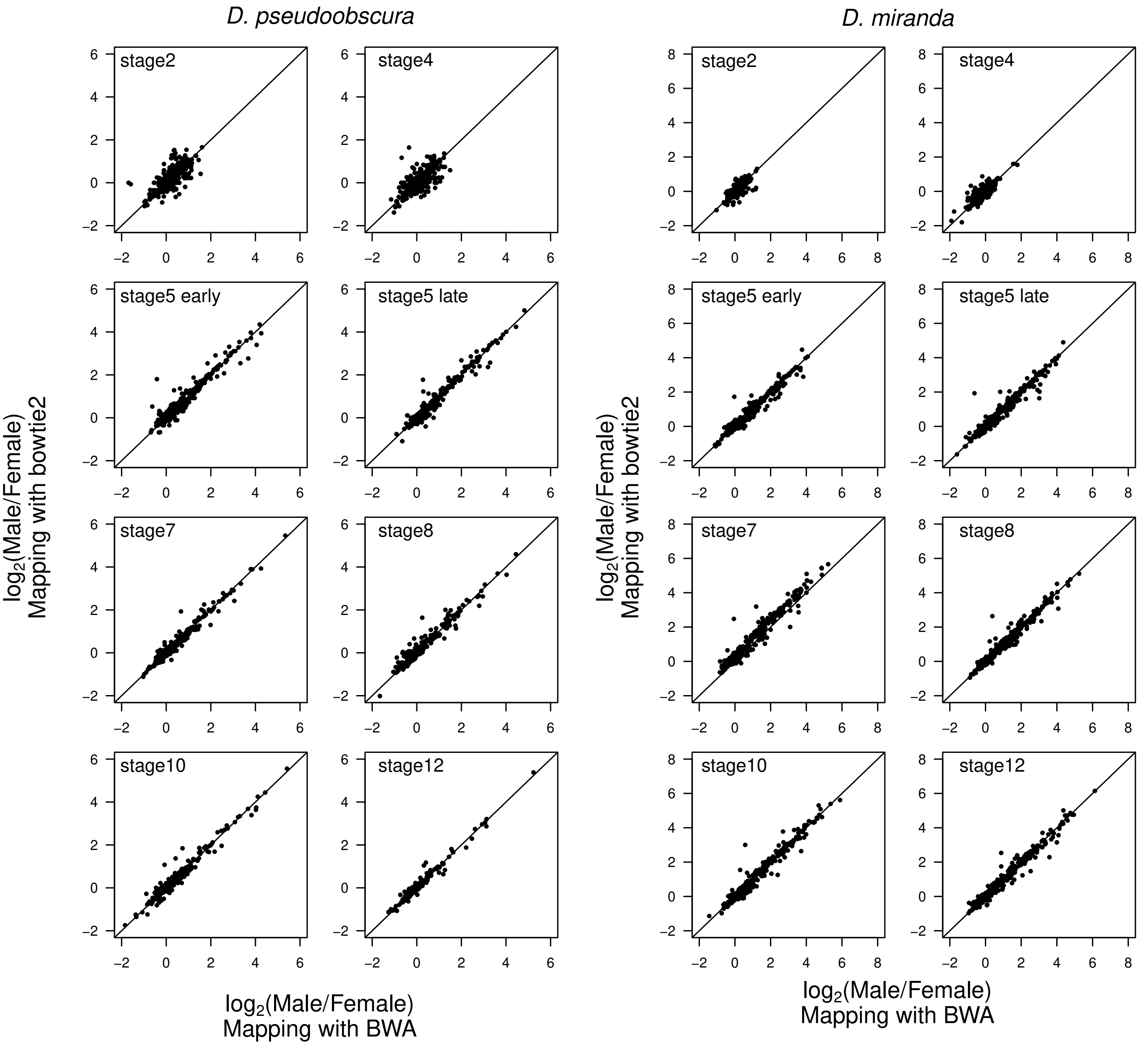


**Supplementary Figure 9. Comparison of read mapping to the TE index using BWA and bowtie2.** The two alignment programs yield highly similar results. The fold differences of TEs between males and females are strongly and significantly correlated accross all samples and stages.

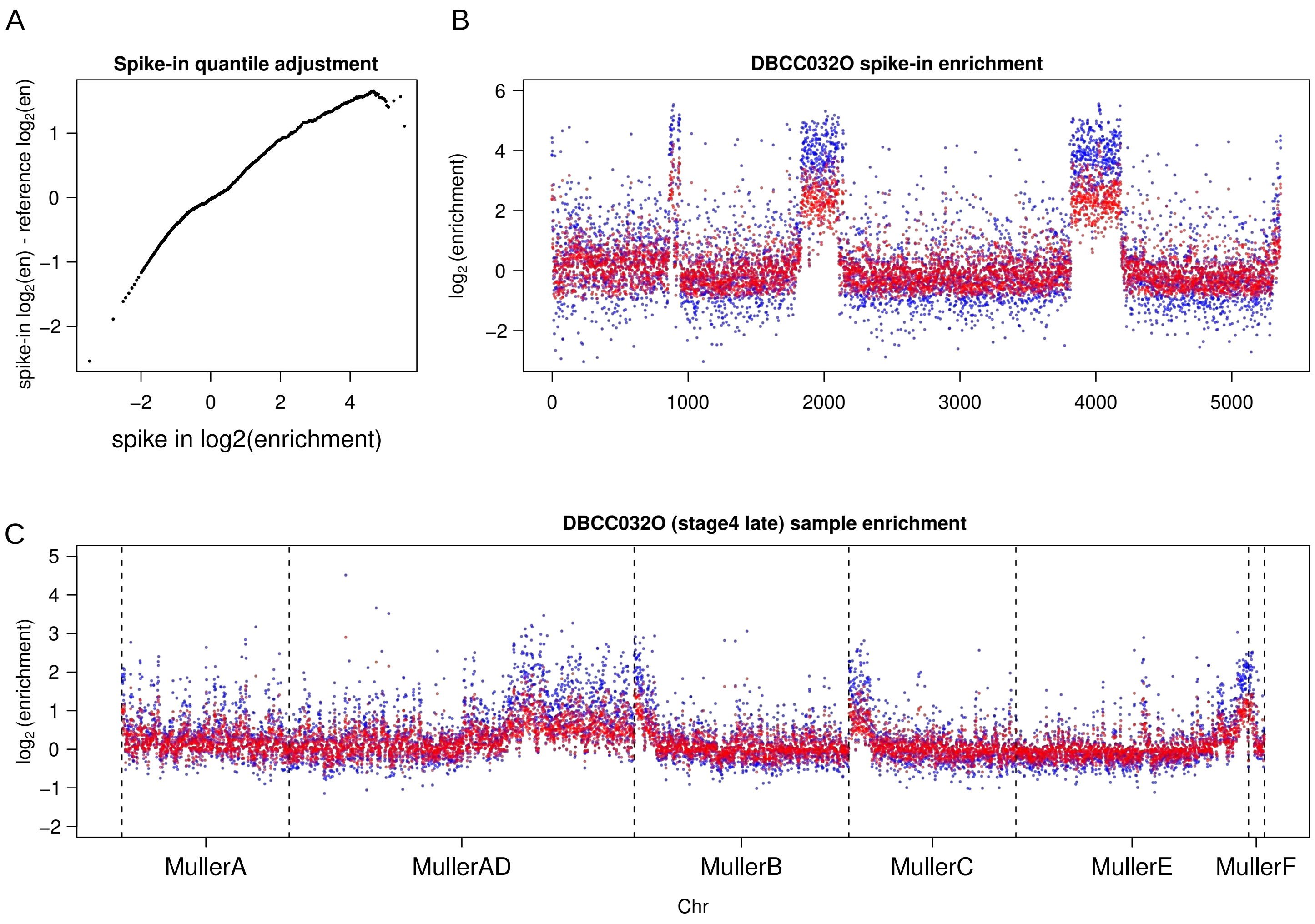


**Supplementary Figure 10. Quantile-informed spike-in normalization procedure.** A. For the spike-in, enrichment (E) at genomic window *i* is first determined by using a simple median autosomal coverage (M) for normalization:

$$E_i = \frac{D_{Ci}/M_C + 0.01}{D_{Ii}/M_I + 0.01}$$

Where C and I are the ChiP and Input samples and the coverage (D). The 0.01 acts as a small pseudocount. To match the quantiles (at 0.1 intervals) of the spike-in with that of the spike-in reference, we subtracted the the log2 enrichment of the former from the latter, generating an adjustment profile. The adjustment profile provides information regarding how much each quantile (dot) and the corresponding enrichment needs to be adjusted to match the spike-in with the reference. B. Based on this adjustment profile (Q), for a given enrichment value (E) across the genome (blue points), the normalized enichment (N) (red points) is then:

$$\log_2(N) = \log_2(E) - Q_E$$

C. The same transformation is then applied to the actual sample. Blue and red points are the enrichment before and after transformation, respectively. Chromosomes are demarcated by dotted lines.
